# Local adaptation in a metapopulation - a multi-habitat perspective

**DOI:** 10.1101/2025.05.03.652039

**Authors:** Oluwafunmilola Olusanya, Nick Barton, Jitka Polechova

## Abstract

This study extends existing soft selection models of local adaptation in metapopulations from two habitats to a multi-habitat scenario, where each habitat exerts unique selection pressures. Specifically, we examine a three-habitat multilocus model in which each allele is favored in habitat 1, disfavored in habitat 3, and the selection pressure in the intermediate habitat may be different across loci. Employing the diffusion and fixed state approximations under the assumption of linkage equilibrium, we investigate conditions for the persistence of a polymorphism. We derive analytical thresholds for such persistence, which reveal scaling for the model parameters, local deme size (N), migration rate(m), selection pressure (*s*_*i*_) and the proportion, (*α*_*i*_) of each habitat. We show that under the assumption of infinitely many islands and selective neutrality in the intermediate habitat, the size of the intermediate habitat does not affect the maintenance of polymorphism. With symmetric selection pressure (*s*_1_=*s*_3_=*s*) in habitats 1 and 3, the system can be fully characterized by the product *Ns*, the product *Nm*, and a parameter *β*, defined as the ratio of the size of habitat 1 (favoring the allele) to habitat 3 (where the allele is disfavored). We find that the range of polymorphism widens as gene flow between demes decreases and the symmetry of habitats increases (*β* approaches 1). In the final section, we explore the effect of drift on the critical migration threshold as well as the effect of symmetry between selection. We demonstrate that genetic drift considerably lowers the critical migration threshold required for the maintenance of polymorphism. Furthermore, when each island is small but there are (infinitely) many of them, relatively low levels of gene flow can have a large impact in preventing genetic differentiation in a fragmented population.

## Introduction

In evolutionary biology, local adaptation is the process by which populations evolve unique traits and genetic variations to better suit their respective habitats (Williams 1996). These adaptations can encompass changes in an organism’s physiology (e.g., metabolic rate), morphology (e.g., body size) or other attributes.

Local adaptation plays an important role in the evolution and survival of populations in heterogeneous environments. It can be a driver for speciation (i..e., the formation of new species) when populations become increasingly specialized to their local conditions and lose their capacity to interbreed with individuals from other populations (Gavrilets 2003). On the contrary, its loss can have serious consequences for species and for the ecosystem they inhabit (Walters and Berger 2019). For example it can lead to increased vulnerability to environmental changes and extinction risks as well as the disruption of ecological interactions (Frankham et al. 2017, Urban 2015) etc. Therefore, understanding the dynamics and factors that promote or constrain such adaptations is crucial for conservation and management efforts. This understanding can help in designing strategies to preserve genetic diversity within metapopulations and further help to predict the response of populations to changing environmental conditions such as climate change and habitat fragmentation.

Local adaptation typically emerges from complex interactions between ecological and evolutionary processes. Ecological processes constitute the interactions between organisms and their environment - when populations are distributed through space (be it as a result of natural or human-induced activities, like habitat fragmentation), they are exposed to different environmental conditions such as variations in abiotic factors like temperature, resource availability, climate, etc., or biotic factors like predation and competition. These environmental differences can cause variation in the selective pressures faced by these populations, promoting the emergence of local adaptations and genetic differentiation among them. A typical example of this are metapopulations of pocket gopher (*Thomomys spp*) in the Great Basin region of North America (Rogers 1991) where populations residing in meadow habitats characterised by depressions (or valleys) with high water availability, have evolved narrower skulls and longer claws for burrowing through soft, moist soils. Whereas, those residing in the sagebrush steppe in more arid regions, have evolved broader skulls and shorter claws to help them traverse drier, more compact soils. These adaptations have enhanced survival and reproduction within each local gopher population.

Evolutionary forces also greatly influence local adaptation. These forces include the strength of natural selection, mutation, recombination, gene flow (due to migration) and genetic drift. Strong selection pressures enhance the spread and accumulation of locally beneficial mutations within sub-populations, increasing fitness as a whole. Mutations generate the novel genetic variants needed for adaptation, and recombination rearranges genetic material, producing novel allelic combinations that can contribute to such adaptations. Migration plays an integral role as it connects demes or sub-populations within a metapopulation. However, it can have discordant consequences (Blanquart et al. 2012, Olusanya et al. 2023, Sachdeva et al. 2022) – it can engender genetic diversity thereby facilitating adaptation through the proliferation of advantageous traits across the metapopulation, yet can nevertheless also lead to the introduction of maladapted alleles to an otherwise perfectly adapted deme, thus disrupting the local gene pool. Such “ migration load” when it become too high, leads to a reduction in fitness thus impeding adaptation and increasing the risk of extinction (Holt and Gomulkiewicz 1997). Finally, genetic drift, the random fluctuation in allele frequency within a population also interferes with local adaptation. This is particularly true in small populations (LaBar and Adami 2017, Whitlock 2000) that are more susceptible to chance events which causes the loss or fixation of particular alleles despite their adaptive value (Blanquart et al. 2012). Understanding how these different forces interact to shape local adaptation is key to elucidating the mechanisms underlying biodiversity and predicting future evolutionary trajectories.

Metapopulation models of local adaptation typically distinguish between two modeling paradigms: hard and soft selection. The hard selection model considers an explicit feedback between ecological processes (in particular, population size) and evolutionary processes (allele frequencies at different loci) in shaping metapopulation dynamics and considers the possibility of both local and global extinction resulting from maladaptation within the metapopulation (Haldane 1956, Szép et al. 2021). In contrast, the soft selection model, which is a useful simplification, ignores this feedback and assumes a constant population size for each subpopulation over time regardless of the level of maladaptation. In this work, we explore the latter model, as this allows us disentangle the dynamics of local adaptation and the maintenance of genetic variation from the confounding effects of demographic fluctuations, thus providing us with a basic understanding of the evolutionary processes at play as well as insights into the stability of metapopulations.

An important concern when exploring these models are the assumptions about the metapopulation landscape. Theoretical models exploring the dynamics of local adaptation with constant deme sizes have mostly focused on the interactions between two niches or habitats (see Maynard (1970), Bulmer (1972), Hoekstra et al. (1985), Barton and Whitlock (1997), Lenormand (2002), Blanquart et al. (2012), Bolnick and Otto (2013), Szép et al. (2021), Barton and Olusanya (2022)). For example, using a two-niche model with migration and opposite selection pressures, Maynard (1970) and Bulmer (1972) showed that for polymorphism to be stably maintained, increasing migration would necessitate a fine balance between niche and selection symmetry as this would ensure that no single allele is overwhelmingly favored in the entire population.

Albeit, since nature is more intricate, many species experience environmental gradients that span more than two distinct environments, which calls for the extension of theoretical models to account for more than two habitats simultaneously. This will not only enable us to better capture the more complex nature of adaptation across a heterogeneous landscape, but will also provide insights into the relative significance of local adaptation versus gene flow in shaping population divergence and maintaining genetic diversity.

In this study, we therefore focus our attention on a metapopulation with more than two habitats while also accounting for the effect of drift. Specifically, we concentrate on simple soft selection models, aiming to identify the key factors or conditions that facilitate the persistence of genetic variation (or polymorphism) in such metapopulations. To achieve this, we rely on mathematical theory based on the diffusion approximation. This approach allows us to describe the dynamics of allele frequencies under soft selection.

## Model and Methods

We consider an infinite-island model (Wright 1931) where the population comprises a large number of demes spread across *n* habitats, each containing a proportion *α*_h_ of demes or niches, such that:

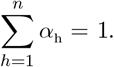

Each habitat (*h*=1, …, *n*) exerts distinct selection pressures, with the fitness of an allele determined by the local environment. Migration occurs between demes at a rate *m*, with individuals entering a common migrant pool before being redistributed uniformly. The model assumes random mating within demes, constant deme size (*N*), and haploid loci with no mutation (so that polymorphism is maintained by other processes like migration and selection).

### Selection Coefficients

Selection pressures differ across habitats, favoring or disfavoring alleles depending on local conditions. For simplicity, we assume fitness is determined multiplicatively across loci, with fitness at any locus *j* proportional to 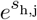.

In habitat *h* = 1, allele *A* is favored (*s*_1,j_ *>* 0), in habitat *h* = 3, allele *A* is disfavored (*s*_3,j_ *<* 0) and in habitat *h* = 2, selection depends on the scenario.

- Scenario 1: Neutral habitat (*s*_2,j_ = 0 for all *j*).
- Scenario 2: Mixed selection (*s*_2,j_ *>* 0 for half the loci and *s*_2,j_ *<* 0 for the other half).

To simplify the model further, we assume linkage equilibrium (LE), disregarding any correlations between loci. This assumption is valid when evolutionary processes occur slowly relative to recombination. Under these assumptions of constant size and LE, allele frequencies at each locus evolve independently and the problem reduces to that of a single locus, where we need only know the average allele frequency across all demes at a given locus.

### Diffusion approximation

Population genetics relies extensively on the diffusion approximation, which establishes a framework for understanding the distribution of allele frequency across a range of models that are equivalent if *s*, 1*/N* ≪1.

In continuous-time and assuming linkage equilibrium, the rate of change in frequency of the *A* allele due to selection, migration and drift at any haploid locus, and in any deme, *i*, in habitat *h* can be written as,

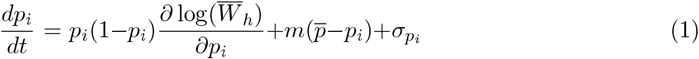

where the direction of evolution is determined by the slope of 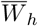, the mean fitness of the habitat in question. 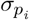 symbolizes the effect of drift and is a real-valued stochastic process with zero mean and covariance 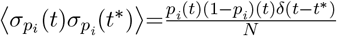. Finally, 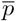 is the frequency of the *A* allele averaged across all demes of the population, is equal to the average in the migrant pool (since migration is uniform), and depends both on the mean frequency of the *A* allele in the different habitats as well as the proportion of demes in these habitats. In other words,

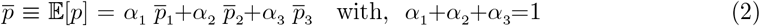

According to Wright (1937) and Kimura (1955), the stationary distribution of allele frequency *ψ*(*p*) in any *i* can be written as,

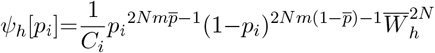

where *C*_*i*_ is the normalization constant. Using the above, the stationary distribution in the three habitats (i.e., *ψ*_1_[*p*_*i*_], *ψ*_2_[*p*_*i*_] and *ψ*_3_[*p*_*i*_]) can be obtained by substituting in *h* and the corresponding 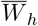 and this will depend on the parameters *Nm* and *Ns*_*h*_. One can then numerically integrate over these distributions to obtain the expectations, 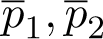 and 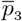, which can now be substituted into eq. (2) to get the mean frequency of the *A* allele in the metapopulation (using a self-consistent iterative process).

### Fixed-state approximation (limit of low migration - *Nm<<*1)

The fixed state approximation is a simplification which assumes that gene flow is limited (i.e., *Nm*≪1) among the different habitats (see Barton and Olusanya, 2022) so that any deme is “ nearly fixed” for one or other allele, with stochastic transitions between fixation for alternative alleles. Under soft selection, this allows us to characterize the genetic state of habitat *h* by the rate of transition (i.e., the rate at which one allele replaces the other in the population) from *a* to *A* (or *A* to *a*). The transition rate from *a* to *A* (*A* to *a*) is simply the product of the fixation probability of the *A* (*a*) allele times the number of new *A* (*a*) alleles entering the population (see equation 2.2 of Barton and Olusanya (2022) where 1≡*A* and 0≡*A* here). Using this approximation, one can then estimate the equilibrium expectation of *A* (*a*) in *h* by the equilibrium proportion of demes fixed for *A* (*a*).

As in Barton and Olusanya (2022), suppose we represent the proportion of demes fixed for the *A* allele in habitat *h* by *P* _*h*_ and that fixed for the *a* allele by *Q*_*h*_ then, focusing on the *A* allele, the evolution of *P* _*h*_ in any *h* can be expressed as,

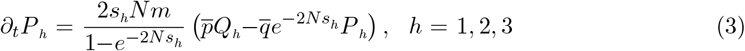

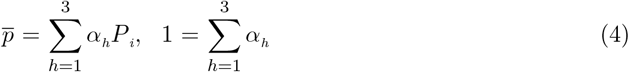

This will be later used to analyze the conditions for a polymorphism.

## Results

### Scenario 1

We begin by considering the case where the *A* allele is favored in habitat 1 (i.e., *s*_1,j_ *>* 0), neutral in habitat 2 (i.e., *s*_2,j_ = 0) and disfavored in habitat 3 (i.e., *s*_3,j_ *<* 0). Since alleles evolve independently, we drop the *j* subscript and focus on the dynamics at a single locus taking *s*_1_ = 1, *s*_2_ = 0 and *s*_3_ = −1 respectively. Our interest is in determining the conditions that favour the persistence of a polymorphism under such a scenario. In particular, we derive critical selection and migration thresholds that allows persistence. But first, let us explore the role of gene flow.

### Role of gene flow on equilibrium allele frequency

Figure 1 shows how the expected frequency of the *A* allele in habitats 1, 2 and 3 respectively (i.e., 𝔼_1_[*p*], 𝔼_2_[*p*] and 𝔼_3_[*p*]) depend on the average allele frequency, *p*, in the migrant pool given different levels of gene flow, *Nm*. The results are obtained numerically using the diffusion approximation.

**Figure 1:**
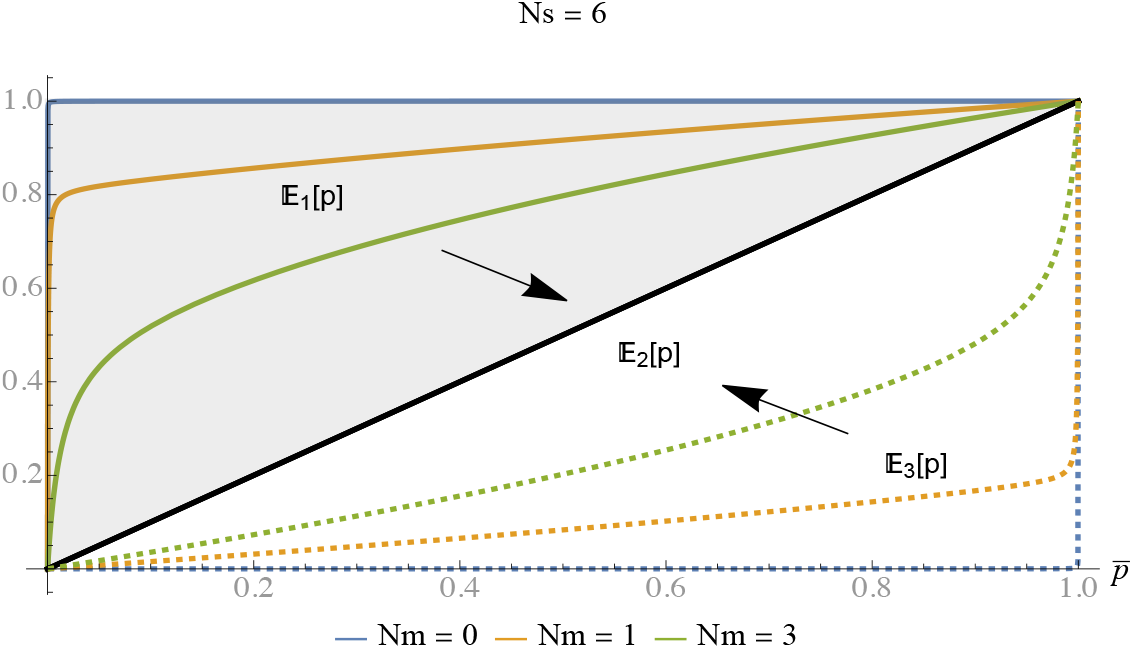
Dependence of the expected frequencies of the *A* allele in the three habitats on 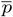 for different levels of gene flow (*Nm*) and with fixed *Ns*. Black arrows pointing towards 𝔼_2_[*p*] indicate that gene flow pushes the allele frequencies in the extreme habitats (i.e., in *h* = 1 and *h* = 3) towards 𝔼_2_[*p*] which coincides with the average, 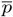 in the migrant pool.

In the absence of gene flow (i.e., *Nm* = 0), the extreme habitats are fully well adapted to their local environmental conditions with the frequency of the *A* allele being 1 in *h*=1 (solid blue line) and 0 in *h*=3 (dashed blue line; so that the *a* allele has frequency 1 here). Increasing gene flow however introduces maladaptation, reducing the frequency of the favored allele in both habitats and pushing them towards the average in the migrant pool.

### Deterministic equilibria

To obtain the equilibria in eq. (2), we numerically plot 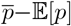 against 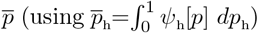 Equilibria correspond to the points where the curves intersect the *x*-axis (see Figures. 2(a) and 2(c)).

**Figure 2:**
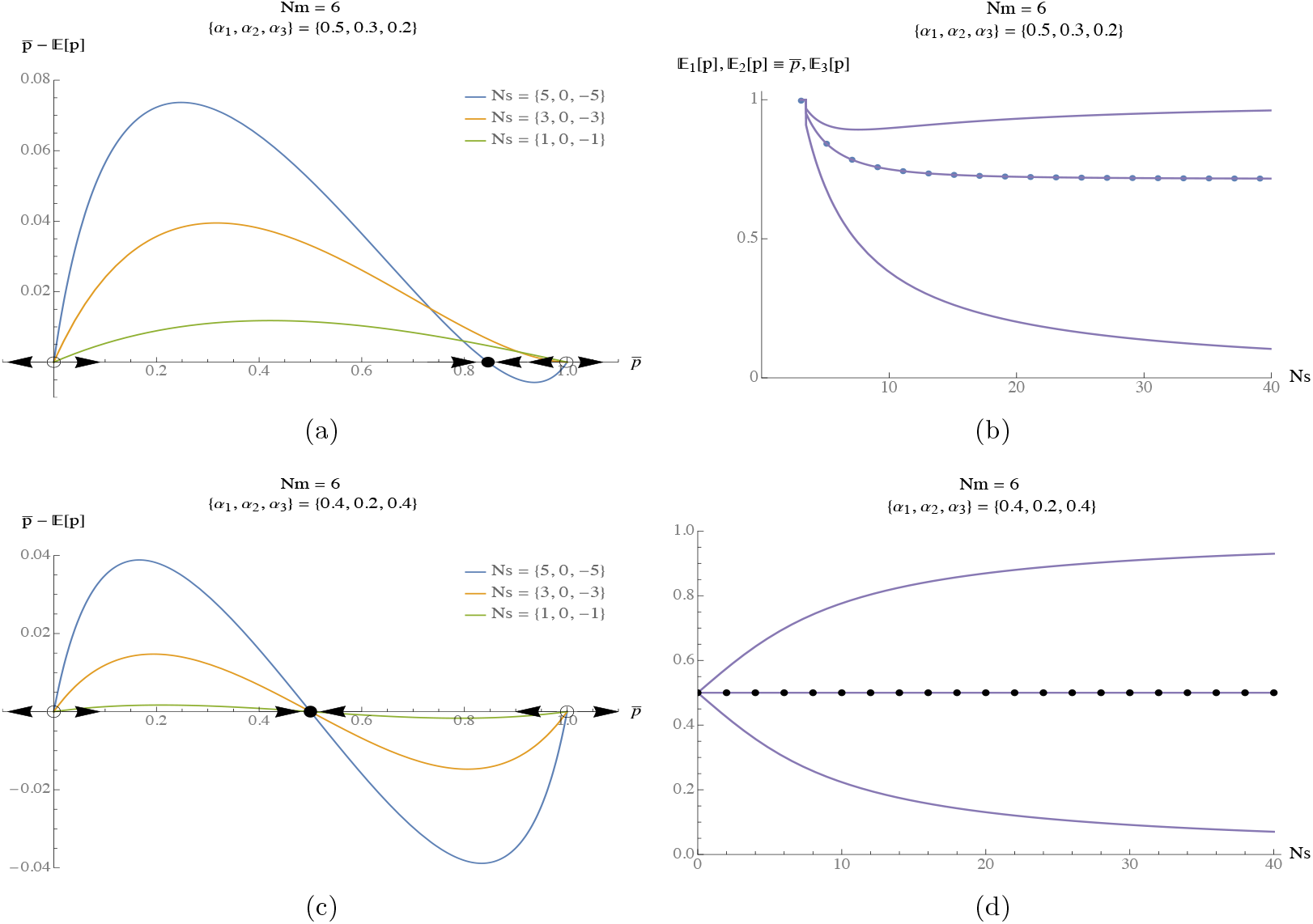
(a.) The three possible equilibria 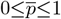 for the *A* allele for a given combination *{α*_1_, *α*_2_, *α*_3_*}* and *Nm* = 6. (b.) A stable polymorphism 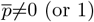 only exists past a critical threshold, *Ns*_*cr*_ = 3.1 for *Nm* =6 and *{α*_1_, *α*_2_, *α*_3_*}* = *{*0.5, 0.3, 0.2*}* (c.)-(d.) With exact symmetry of *α*_1_ and *α*_3_, the stable polymorphism always exists at 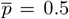 independent of *Nm* and *Ns*. Black dots in figs. (b.) and (d.) represent the average allele frequency 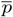 across the metapopulation. Results are obtained numerically using the diffusion approximation.

We find (Figure 2(a)) that a stable polymorphism 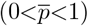 always exists for any combination of *α*_1_, *α*_2_, *α*_3_ provided that selection is strong relative to migration (i.e., *Ns*≳*Nm*, blue line) as can also be seen in Figure 2(b) (black dots). This means that if *Ns* is less than a given critical threshold value, which we will denote by *Ns*_cr_ (in this example *Ns*_cr_=3.1, Figure 2(b)), polymorphism will be lost and one of the alleles (in this case the *A* allele) would fix throughout the metapopulation (see lhs of Figure 2(b)).

A trivial case occurs when the two extreme habitats (*α*_1_ and *α*_3_) are precisely balanced (i.e., are equally common or rare), then there would always exist a stable polymorphism at *p* = 0.5 independent of *Nm* and for all values of *Ns* (as can also be seen in Figure 2(d)).

In this study, we are interested in cases where 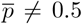 as this provides a basis for exploring critical thresholds. In particular, we consider situations where *α*_1_*>α*_3_ independent of *α*_2_ (i.e., the habitat where the *A* allele is favored has a larger proportion of demes compared to the habitat where it is disfavored).

### Maintenance of polymorphism and critical thresholds

Here, we consider how the persistence of a polymorphism is influenced by factors such as migration, selection, demic proportion and drift as well as provide an analytical handle on thresholds for persistence.

Figure 3(a) shows that when selection is weak (or comparable) relative to migration, allele *A* eventually fixes across the metapopulation (hence an initial increase in load Figure 3(b)) and this load decreases as the intensity of selection increases. Figure 3(c) further shows that limited migration favors the maintainance of polymorphism due to the lower homogenizing (and hence deleterious) effect of migration whereas, increasing *Nm* causes the *A* allele to invade and fixe across the metapopulation. Interestingly, all that matters here is the ratio of the proportion of demes where the *A* allele is favored to that where it is maladaptive i.e., *α*_1_*/α*_3_, and not the actual proportions *α*_1_ and *α*_3_. We call this ratio *β* (i.e., *β*:=*α*_1_*/α*_3_). The lower the value of *β*, the longer the polymorphism persists (i.e., the wider the range of polymophism possible), despite increasing *Nm*, and thus, the higher the critical migration threshold below which polymorphism is possible. This makes sense, as the potential for swamping increases the more dissimilar the extreme habitats are.

**Figure 3:**
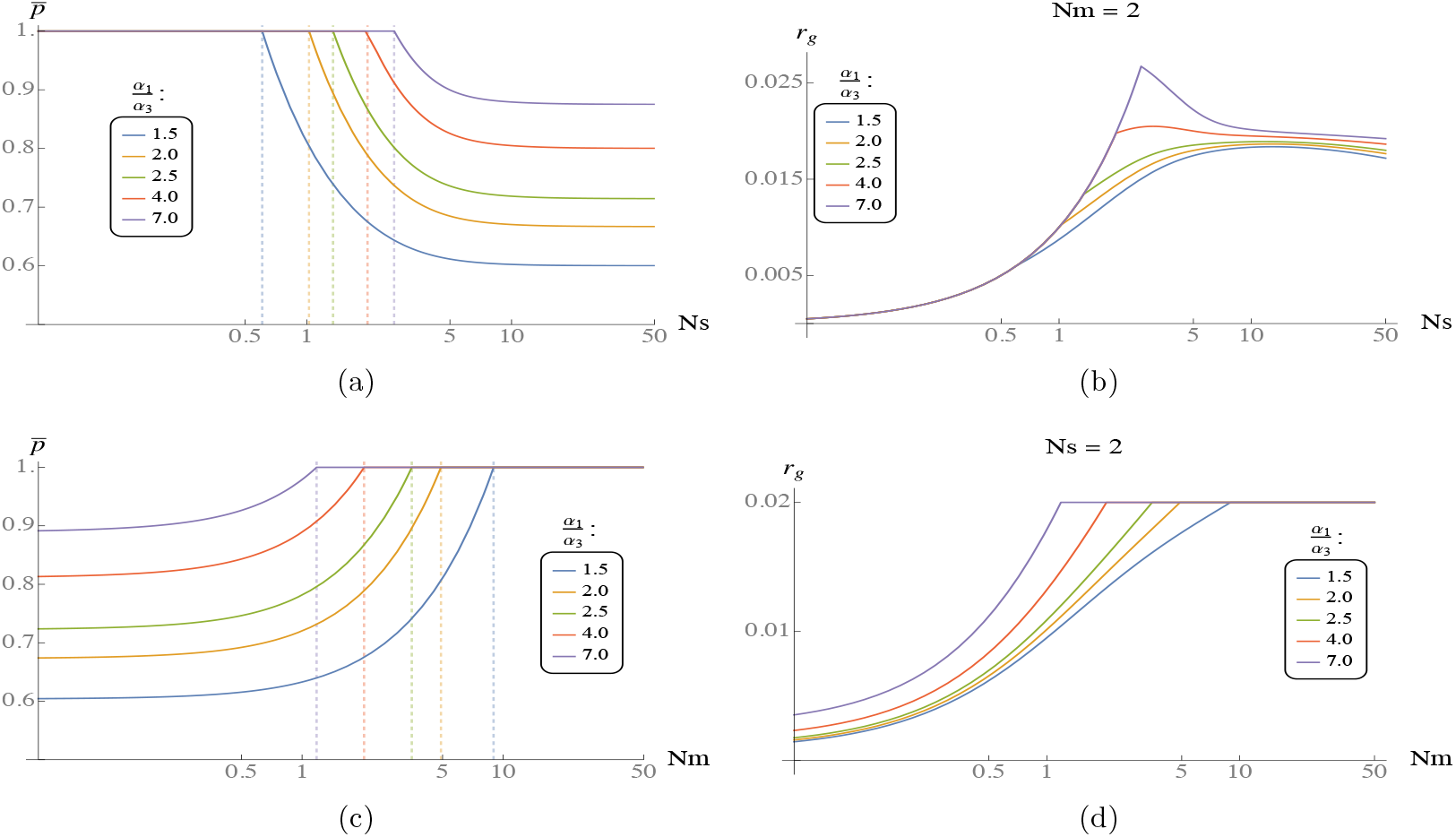
Dependence of the equilibrium average allele frequency on *Ns* (a.) and *Nm* (c.). Dashed lines in (a.) represent the critical selection thresholds, *Ns*_*cr*_, above which a polymorphisn is possible and those in (c.) represent the critical migration threshold, *Nm*_*cr*_ below which a polymorphism is possible. (b.) and (d.) show the mean load in the metapopulation and how this depends on *Ns* and *Nm* respectively. The *x*-axis is plotted on a log scale to better visualise behaviour at longer ranges. Results are obtained numerically using the diffusion approximation.

### Scenario 2

We now investigate whether or not there is an advantage to having an intermediate habitat where half of the loci in this habitat favors the *A* allele, whereas, it is disfavored at the remaining half. Does this for instance make it easier to maintain polymorphism? In other words, we consider a scenario where {*s*_*2*,*1*_, …, *s*_*2*,*j*_}={−1, −1, …, +1, +1}. However since loci are decoupled under our model, we can simply follow the dynamics at a single locus conditioned on 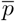 at that locus. Hence, considering the dynamics at locus 1 and dropping the *j* index, we have *s*_1_=+1, *s*_2_=−1 and *s*_3_=−1.

Figure 5(a) (and 5(b)) in appendix A. show that the critical threshold, *Ns*_*cr*_ (and *Nm*_*cr*_) as well as the dynamics past this threshold, i.e., *Ns>Ns*_*cr*_ (and *Nm<Nm*_*cr*_) are exactly the same as can be seen in fig. 3(a) (and 3(c)) provided that *β* is the same in both scenarios (compare blue and red curves in fig. 5(a) (see appendix A). and fig. 3(a) as well as in 5(b) (appendix A). and 3(c)). In fig. 3(a) (blue curve), the actual demic combination used to obtain the plot was {*α*_1_, *α*_2_, *α*_3_}={0.3, 0.5, 0.2} meaning that the *A* allele was favored in 30% of demes, selectively neutral in 50% of demes and disfavored in 20% of demes so that *β*=1.5. However, in fig. 5(a), the combination used was {*α*_1_, *α*_2_, *α*_3_}={0.6, 0.2, 0.2} meaning that the *A* allele was favored in 60% of demes and disfavored in 40% of demes so that *β* is again 1.5. In essence, we see that what really matters for the overall dynamics is not the individual proportion of demes but the value of *β*. So for a single locus, independent of whether selection is neutral or disadvantageous in the intermediate habitat, we will obtain similar dynamics overall provided that *β* is the same.

In fact, for a single locus, this second scenario considered above, i.e., with *s*_1_=1, *s*_2_=−1 and *s*_3_=−1 reduces to the two habitat case considered in Szép et al. (2021) where we only need to know the proportion of demes where the allele is favored i.e., *α*_1_ (since the proportion where it is disfavored can be obtained simply as 1−*α*_1_).

The trivial behaviour observed above can be attributed to the assumptions of our model. With hard selection however (not assumed here), where alleles co-evolve with each other and are coupled via *N*, we would expect to see a non-trivial dynamics.

Next, we obtain analytical formulas for critical thresholds (focusing strictly on the case *s*_1_=1, *s*_2_=0, and *s*_3_=−1). To do this, we employ the fixed state approximation. First, we focus on the critical selection threshold, *Ns*_*cr*_, above which a polymorphism is possible. This can be split into threshold values when gene flow among the different habitats is limited (*Nm*≪1) and when it is abundant (*Nm*≫1). In the limit of low migration where allele frequency distribution are bimodal with loci nearly fixed for the *A* or *a* allele, the equilibrium mean allele frequency 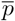 across the metapopulation can be obtained by first setting the lhs of eq. (3) to 0 and solving for *P*_1_, *P*_2_ and *P*_3_ respectively. These can now be substituted into eq. (4) to obtain 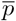 as,

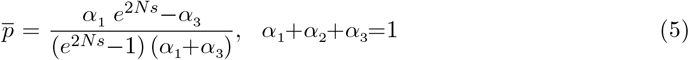

Equation (5) can then be solved for *Ns* yielding 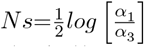. So, in the limit of low *Nm*, we require a selection strength above ((1*/*2*N*) *log*(*α*_1_*/α*_3_)) to maintain a polymorphism. To find the threshold value for larger values of *Nm*, we use a deterministic analysis (see also soft selection analysis in Szép et al. (2021)). Just above the critical selection threshold, 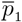 will be close to 1, 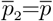 and the difference between 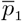 and 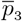 will be very slight (see Figure 2(b) for example). Thus, we can set 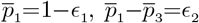 and 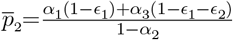. Substituting these into eq. (1), replacing *p*_*i*_ with 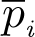 (where *i*=1, 2, 3) and setting 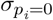 (since we’re dealing with a deterministic analysis), we obtain differential equations, 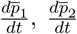 and 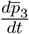 respectively. Consequently, solving 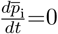 for 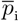 (retaining only lower order terms) we obtain the deterministic threshold as 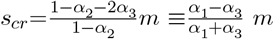. Hence, we have,

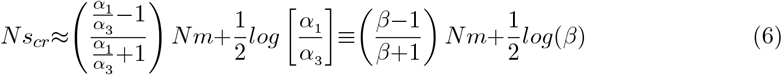

In a similar fashion, using eq. (6), we obtain an analytical expression for the critical migration threshold, *Nm*_*cr*_ as,

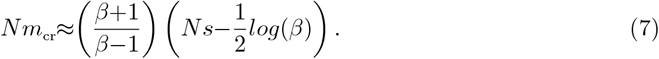

Equations (6) and (7) are approximate equations obtained by adding together the *Nm*→0 (i.e., fixed state) threshold and the deterministic threshold.

Interestingly, both equations (i.e., eq. (6) and (7)) depend only on *β* (*i*.*e*., *α*_1_*/α*_3_) and not on *α*_2_ implying again that in this case, having an intermediate habitat makes no difference to critical selection and migration thresholds.

A comparison of these two equations (i.e., eq. (6) and (7)) with numerically obtained values from the diffusion approximation (see Figure 8(b) in appendix D.) shows a close fit between our derived formulae and the numerical expectation. In particular, as the rate of gene flow (*Nm*) increases (Figure 8(a) in appendix D.), we observe a corresponding rise in *Ns*_*cr*_ (the critical selection threshold) due to heightened migration load within the population resulting from increased gene flow. Consequently, stronger selection is necessary to counteract this effect. *Ns*_*cr*_ is also higher with higher *β*:=*α*_1_*/α*_3_ allowing for polymorphism in a restricted range of selection intensity when the two extreme habitats are more dissimilar in proportion.

So far, we have established that in the symmetric case (*s*_1_=*s*_3_=*s*), the intermediate habitat (i.e., where *s*_2_=0) makes no difference to critical migration and selection thresholds (see also eq. (6) and (7)). To check whether this habitat matters in any other way (i.e., past critical thresholds), we compare the dynamics of 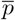 past the threshold values i.e., at *Nm<Nm*_*cr*_ (*Ns>Ns*_*cr*_) for two metapopulations with equal *β* and different *α*_2_. In particular, for the two metapopulations, we use the demic proportions {0.2, 0.7, 0.1} and {0.4, 0.4, 0.2} respectively. Our results, Figure 6(a) in appendix B. (and Figure 7), show similar dynamics (divergence) for *p* for *Nm<Nm*_*cr*_ (and *Ns>Ns*_*cr*_) for both metapopulations suggesting the independence of these results on *α*_2_. We furthermore check if this conclusion holds true with asymmetric selection (i.e., with *s*_1_ ≠*s*_3_) and find that even under the assumption of asymmetry, *α*_2_ has no influence on critical migration thresholds for polymophism or on the divergence of 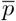 at *Nm<Nm*_*cr*_ (see Figure 6(b) in appendix B.).

Finally, we quantify the effect of drift (finite deme sizes) on the maintenance of polymorphism and how this depends on habitat proportions (Figures 4(a) and 4(b)) while also relaxing the assumption of symmetric selection. To do this, we consider the *s*_1_, *s*_3_ region within which a polymorphism persists and explore its dependence on different deme sizes *N* (starting from larger *N* to lower *N*). Although what really matters for polymorphism are the scaled parameters *Ns*_1_, *Ns*_3_ and *Nm*, plotting this way allows us to easily draw comparison and identify whether for a given *s*_1_, *s*_3_ value, polymorphism is better maintatined in a metapopulation with larger or smaller deme sizes.

**Figure 4:**
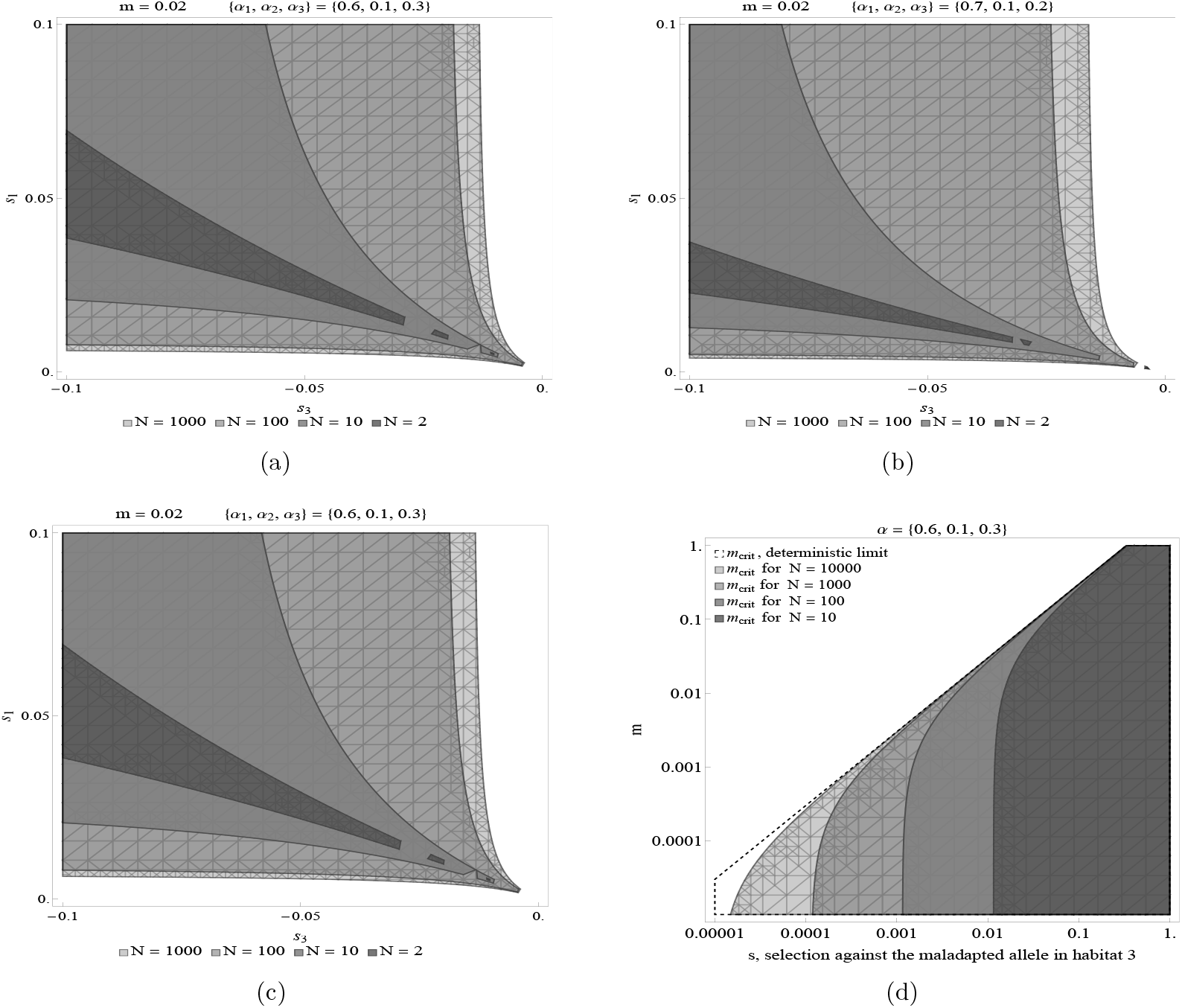
(a.)-(b.) Effect of drift on *s*_1_*/s*_3_ region for which polymorphism is possible (*s*_1_ and *s*_3_ are the strength of selection for and against the *A* allele in habitats 1 and 3 respectively). We have also relaxed the assumption of symmetric selection here. Contrasting (c.) with (a.) show that even with drift, the intermemdiate habitat play no role in the maintenance of a polymorphism provided that *β* is constant (*α*_2_=0.1 in (a.) and 0.4 in (c.) and *β*=2 in both plots). (d.) Effect of drift on the critical migration threshold below which polymorphism is possible. (a).-(d.) are obtained from the diffusion approximation. (c.) is obtained using eq. (7).

We see from Figures 4(a) and 4(b) that increasing drift (i.e., increasing 1*/N*) reduces the overall genetic diversity in the metapopulation and constrains the region for which polymorphism is possible. This is because with increasing drift, certain alleles become more or less common purely by chance, leading to a decrease in the overall number of the two alleles segregating in the population. This reduction in genetic diversity consequently limits the potential for polymorphism. We also observe that this behaviour is more pronounced with higher *β* (i.e., with *α*_1_≫*α*_3_). A comparison between Figures 4(a) and 4(c) indicates that even with drift, the intermediate habitat does not matter for the maintenance of a polymorphism (as seen in the observed independence of the results on *α*_2_ in both plots) provided that *β* remains constant.

Furthermore, we explore the effect of drift on the critical migration threshold necessary for a polymorphism. As illustrated in fig. 4(d), genetic drift considerably lowers the critical migration threshold required for a polymorphism so that relatively low levels of gene flow can have a large impact in preventing genetic differentiation resulting from drift.

## Discussion

The preservation of genetic polymorphisms within metapopulations has been a topic of significant interest in population genetics. Our work builds upon previous research on local adaptation in a metapopulation involving two habitats, extending the analysis to three-habitats. Our findings offer insights into the interplay between selection, migration, drift and habitat proportions in maintaining genetic diversity within metapopulations.

One key finding of our study is the notable increase in the range of polymorphism with limited migration between demes. This is consistent with previous research (albeit involving two habitats) where reduced gene flow is found to enhance local adaptation and promote genetic diversity within metapopulations. Constrained gene flow fosters the persistence of polymorphism by preventing the rapid homogenization of alleles across habitats. This reinforces the fact that in nature, factors such as dispersal barriers or other mechanisms that hinder gene flow could be vital in preserving genetic diversity within metapopulations. For a given *Nm*, we also find that the range of polymorphism is further increased if the habitat where the allele is well adapted and maladaptive respectively are in roughly equal proportions. This suggests that when selective pressures in these habitats are evenly matched, this creates an environment where polymorphism can thrive. This may arise from a dynamic equilibrium in which opposing selection strengths sustain a stable polymorphic state, thereby preventing the complete fixation of one allele over the other.

Our study also highlights the crucial role of selection relative to migration (*Ns*≳*Nm*) in driving the persistence of polymorphism. Strong selection counteracts the homogenizing effect of gene flow ensuring the coexistence of both alleles in the metapopulation. Conversely, genetic drift constrains the region within which polymorphism is possible.

Under our (single locus) model of soft selection, we found no clear advantage for the maintenance of polymorphism when there is an intermediate habitat where alleles are selectively neutral and this holds true away from critical thresholds (i.e., for *Ns>Ns*_*cr*_ and *Nm<Nm*_*cr*_) and with asymmetric selection (i.e., with *s*_*1*,*j*_ *≠s*_*3*,*j*_). Instead, our findings emphasize that what really matters for the persistence of polymorphism is the relative balance of favorable and maladaptive habitats. Specifically, the ratio of the proportion of demes where the allele is favored to where it is maladaptive (i.e., *β*) emerges as an important parameter driving the dynamics of polymorphism. This result however holds under the infinite island model assumed in this work. With finite islands, genetic variation (and hence polymorphism) could be lost by chance (see Barton and Olusanya 2022) and a low amount of mutation may therefore be required to maintain variation in the long term.

Finally, we derived analytical formulas for the critical selection and migration thresholds for polymorphism in the three-habitat case. These formulas can provide useful insights into the parameter ranges where a polymorphism is possible.

While this study represents a step forward in understanding local adaptation and the maintenance of polymorphism in metapopulations with more than two habitats, several avenues for future research remain open. Investigating the dynamics of adaptation under a model of hard selection (where we account for changes in population size via allele frequency changes and vice versa i.e., explicit eco-evo feedback) are promising directions for further exploration. In this case having an intermediate habitat where the A allele is at a selective advantage at half the loci and at a disadvantage at the remaining half (or any other complex architecture) could produce interesting dynamics and reveal novel insights into local adaptation and the maintenance of polymorphism.

Secondly, relaxing the assumption of linkage equilibrium and exploring the role of non-random associations or interference between loci could provide a more realistic depiction of genetic interactions within metapopulations. In this case, the net effect of other loci on any selected locus can be captured using the effective migration approximation^1^ (see Sachdeva 2022)).

Another compelling avenue for future exploration would be to consider explicit spatial structure. Incorporating explicit spatial configuration or arrangement of populations into our framework may enable a more precise investigation into the role of the parameter *β* for the maintenance of variation and how this is influenced by habitat connectivity. It may also provide a more nuanced understanding of how gene flow, genetic drift and spatial heterogeneity influences the stability and maintenance of polymorphism, offering a more realistic perspective on the mechanisms that shape the genetic composition of natural populations.

Finally, exploring the impact of various ecological factors such as variation in carrying capacities across the different habitats could offer a richer understanding of the interplay between genetic and ecological dynamics in shaping and sustaining genetic diversity within fragmented landscapes.

## Appendices

### A. Maintenance of polymorphism and critical thresholds for the case *X*_1_=1, *X*_2_=−1, and *X*_3_=−1

**Figure 5:**
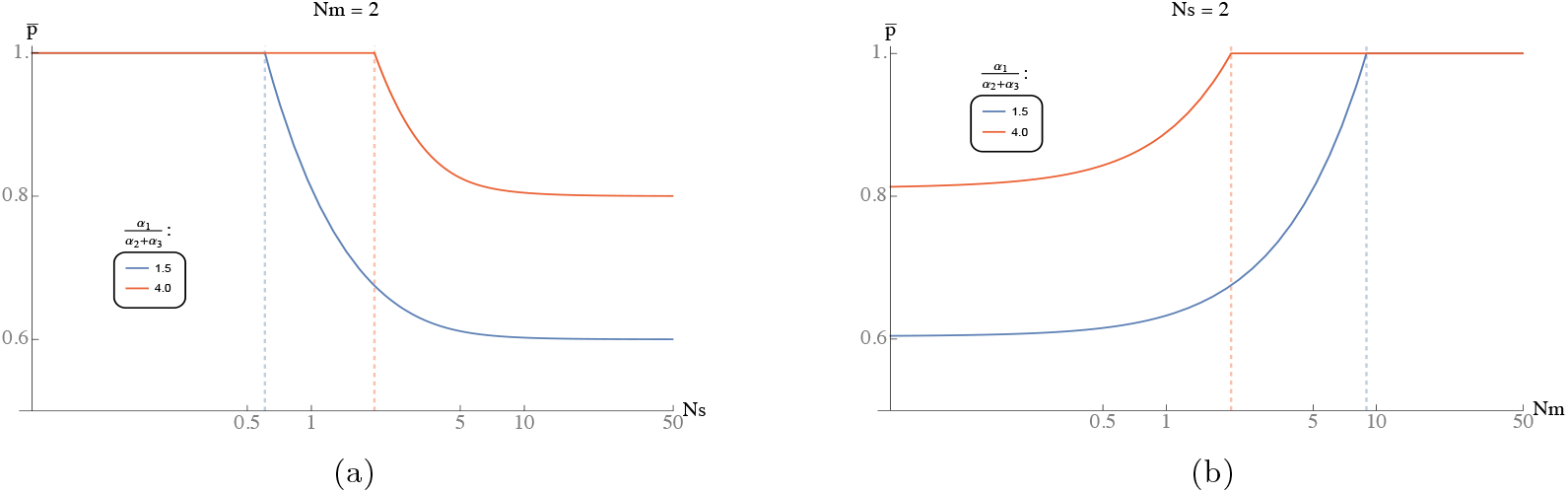
Critical: (a.) *Ns* threshold above which and (b.) *Nm* threshold below which a polymorphism is possible.

### B. Critical migration threshold for polymorphism with similar *β* but different *α*_2_ values. *X*_1_=1, *X*_2_=0, *X*_3_=−1

**Figure 6:**
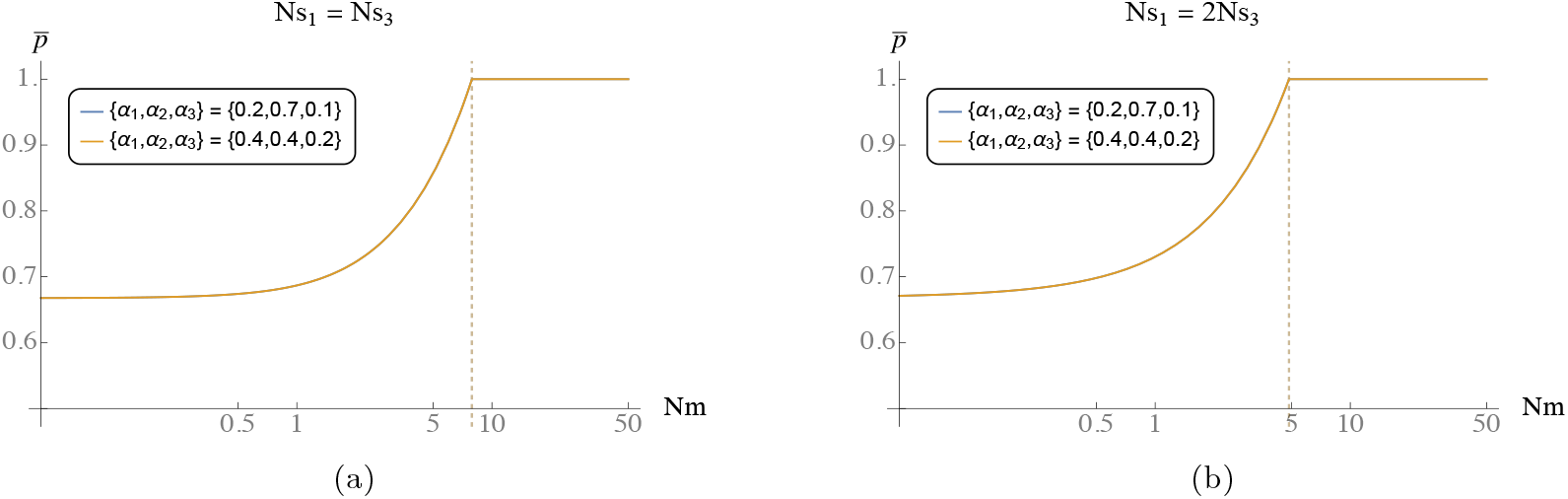
(a.) Symmetric selection *Ns*_1_=*Ns*_3_ (in magnitude). (b.) Asymmetric selectiom.

### C. Critical selection threshold for polymorphism with similar *β* but different *α*_2_ values. *X*_1_=1, *X*_2_=0, *X*_3_=−1

**Figure 7.**
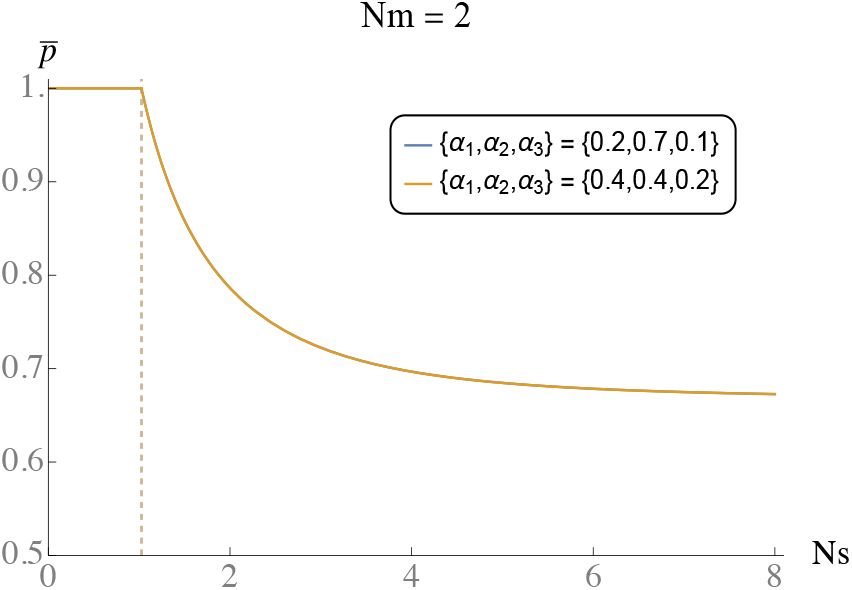

### D. Comparing *Ns*_*cr*_ and *Nm*_*cr*_ with numerical solution from the diffusion approximation

**Figure 8:**
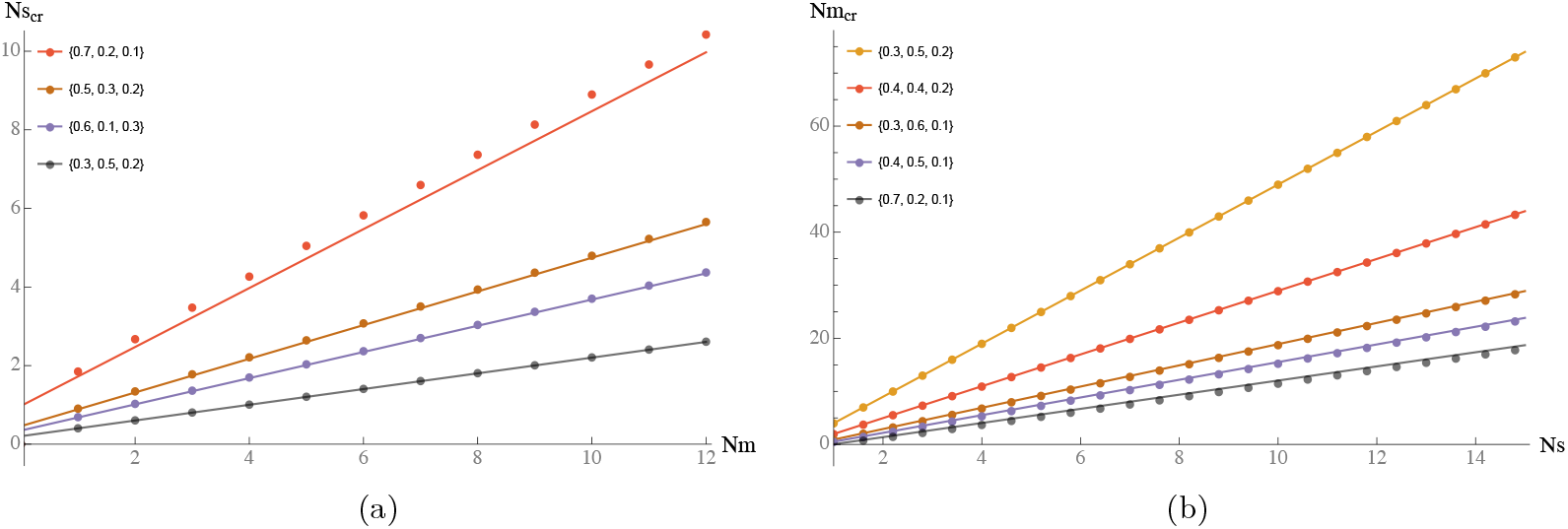
Critical: (a.) *Ns* threshold above which and (b.) *Nm* threshold below which a polymorphism is possible. The different colors represent different *{α*_1_, *α*_2_, *α*_3_*}* combinations. Dotted lines are numerical solutions from the diffusion approximation and solid lines results from eq. (6).

### E. Effect of drift on the maintenance of a polymorphism

**Figure 9:**
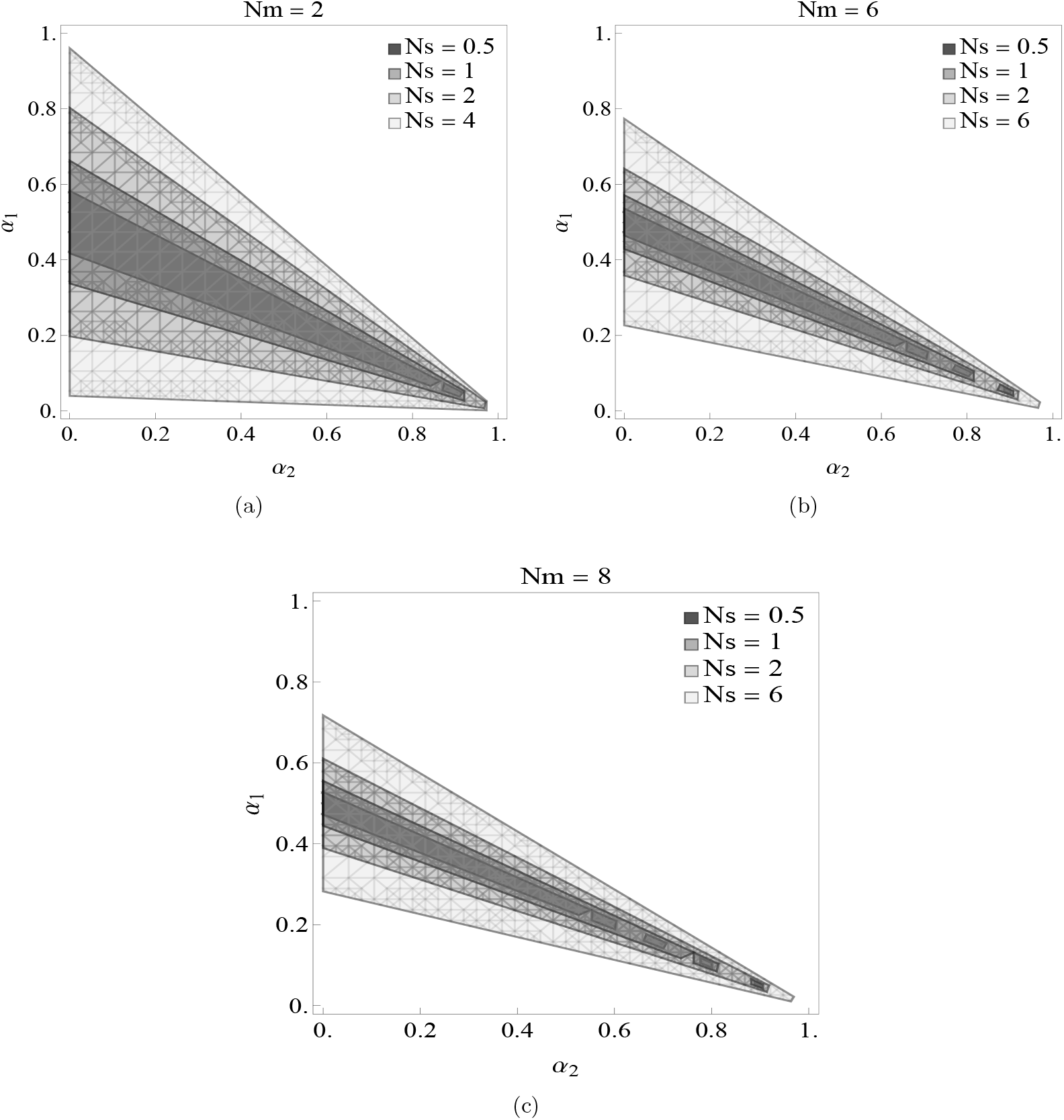
Drift constrains the region within which a polymorphism is possible.

## Footnotes

1 The effective migration approximation accounts for how multilocus LD influence allele frequencies by assuming that it essentially alters the effective migration rate of deleterious alleles, so that the allele frequency distribution at any given locus can still be obtained using the diffusion approximation with the the rate of migration, *m* replaced by an effective migration rate, *m*_*e*_.

